# The effect of phosphatidylserine on the membrane insertion of the cancer-targeting ATRAM peptide is defined by the non-inserting end

**DOI:** 10.1101/623967

**Authors:** V. P. Nguyen, A. C. Dixson, F. N. Barrera

**Affiliations:** University of Tennessee

## Abstract

The acidity-triggered rational membrane (ATRAM) peptide was designed to target acidic diseases such as cancer. An acidic extracellular medium, such as that found in aggressive tumors, drives the protonation of the glutamic acids in ATRAM, leading to the membrane translocation of its C-terminus and the formation of a transmembrane helix. Compared to healthy cells, cancerous cells often increase exposure of the negatively charged phosphatidylserine (PS) on the outer leaflet of the plasma membrane. Here we use a reconstituted vesicle system to explore how phosphatidylserine influences the interaction of ATRAM with membranes. To explore this, we used two new variants of ATRAM, termed K2-ATRAM and Y-ATRAM, with small modifications at the non-inserting N-terminus. We observed that the effect of PS on the membrane insertion p*K* and lipid partitioning hinged on the sequence of the non-inserting end. Our data additionally indicate that the effect of PS on the insertion p*K* does not merely depend on electrostatics, but it is multifactorial. Here we show how small sequence changes can impact the interaction of a peptide with membranes of mixed lipid composition. These data illustrate how model studies using neutral bilayers, which do not mimic the negative charge found in the plasma membrane of cancer cells, may fail to capture important aspects of the interaction of anticancer peptides with tumor cells. This information can guide the design of therapeutic peptides that target the acidic environments of different diseased tissues.

**Statement of Significance:** Current targeted therapies for cancer have limited success due to drug resistance. Resistance often arises after mutation of the receptor being targeted. A more general target is needed to prevent drug resistance. Most aggressive solid tumors have an extracellular medium. We propose that extracellular acidity is promising for improved targeted therapies. The acidity-triggered rational membrane (ATRAM) inserts in membranes only under acidic conditions. However, it is now known how the lipid changes that occur in the plasma membrane of cancer cells impact the membrane insertion of ATRAM. Here we perform biophysical experiments that show that PS, a lipid exposed in the cancer cell, can impact the membrane insertion of ATRAM. We also uncovered a region of the peptide important for insertion.

## Introduction

Efficacious targeting of specific molecular markers in cancer is limited by tumor heterogeneity and drug resistance acquired due to rapid mutation ^*1*^. Therefore, it can be advantageous to target instead an intrinsic tumor property, such as extracellular acidity^*2*^. Cancer cells preferentially metabolize glucose in the presence of oxygen due to the Warburg effect, resulting in increased secretion of lactic acid^*3*^. Combined with poor tumor perfusion, this leads to the accumulation of acidic metabolites that reduce the pH of the extracellular matrix^*4*^. We developed the <u>a</u>cidity-triggered <u>ra</u>tional <u>m</u>embrane (ATRAM) peptide to target the acidic extracellular environment of tumors^*5*^. Biophysical and cellular experiments have shown that the membrane interaction of this highly soluble peptide is pH-dependent. At physiological pH, ATRAM partitions to lipid membranes, but at acidic pH it inserts into the membrane as an alpha helix. ATRAM has been able to deliver a cell-impermeable toxin into HeLa cells, as well as efficiently target solid tumors in mice^*6*^. The insertion p*K* of ATRAM in POPC (1-palmitoyl-2-oleoyl-sn-glycero-3-phosphocholine) vesicles was previously determined to be 6.5^*5*^. This pH is at the lower limit of tumor extracellular acidity^*7, 8*^. To address the need to target mildly acidic tumors, here we worked to tune the tumor-targeting properties by designing two variants with N-terminal modifications. The effect of positively charged amino acids and aromatic amino acids on the insertion of ATRAM was studied.

Previous biophysical experiments studying the membrane interaction of ATRAM have been limited to POPC^*5, 6*^. However, the lipid bilayer of the plasma membrane is highly complex, as both leaflets contain a variety of different lipids^*9*^. A particularly important lipid is phosphatidylserine (PS). PS represents approximately 10% of the phospholipids in the plasma membrane, and remains largely sequestered in the inner leaflet of healthy cells due to lipid asymmetry^*10*^. However, cancerous cells can lose this asymmetry and as a result contain PS on both leaflets of their plasma membranes^*11, 12*^. Unlike PC lipids that contain a zwitterionic head group, PS is a net negatively charged phospholipid. ATRAM has a strong negative net charge at physiological pH; therefore, electrostatic repulsion may emerge between the negative charges of the peptide and the membrane of tumor cells, and control peptide-lipid interactions.

Here we show that the membrane insertion p*K* of ATRAM did not change with increasing PS concentrations, but how the peptide interacted with the membrane was altered. In contrast, in the N_t_ variants PS, had a significant effect on membrane interaction and insertion. We investigated electrostatic interactions as a possible explanation for these different behaviors by performing additional experiments in the presence of NaCl. By understanding how negatively charged lipids affect the insertion of ATRAM into membranes, we can rationally fine-tune the sequence of ATRAM to increase its efficacy to insert into cancer cell membranes.

## Methods and Materials

### Preparation of liposomes

Stocks of POPC and POPS (Avanti Polar Lipids, Inc., Alabaster, AL) were prepared in chloroform. Lipids were dried flushing with argon gas and placed under a vacuum overnight. The resulting dried lipid films were hydrated with 10 mM sodium phosphate (pH 8.0) and extruded with a Mini-Extruder (Avanti Polar Lipids, Inc., Alabaster, AL) through 100 nm polycarbonate filters (Whatman, Maidstone, United Kingdom) to create large unilamellar vesicles (LUVs).

### Peptide labeling

The C-terminal Cys of the ATRAM or K2-ATRAM variants (Table 1) was conjugated with the environmentally sensitive dye NBD, [N, N’-dimethyl-N-(iodoacetyl)-N’-(7-nitrobenz-2-oxa-1,3-diazol-4-yl)ethylenediamine] (Thermo Fisher Scientific, Inc., Waltham, MA). Y-ATRAM (Table 1) was labeled at the N-terminus with succinimidyl 6-n-7-nitrobenz-2-oxa-1,3-diazol-4-yl amino hexanoate (NBD-X SE; Anaspec, Fremont, CA). Free NBD dye was removed by gel filtration using a PD-10 column (GE Healthcare Life Sciences, Marlborough, MA), while labeled and unlabeled peptide were separated by reverse-phase HPLC (Agilent, Santa Clara, CA). The purity of the conjugations was confirmed by MALDI-TOF (Bruker, Billerica, MA).

**Table 1.** Sequences of the ATRAM peptide and variants. Residues in red are the N-terminal modifications, the single tryptophan is marked in blue, and the key residue E12 is shown in green. Black G residues in bold are replaced with C for conjugation.

| Peptide | Sequence |
| --- | --- |
| ATRAM | GLAGLAGLLGLEGLLGLPLGLLEGLWLGLELEGN |
| K2-ATRAM | GKAGKAGLLGLEGLLGLPLGLLEGLWLGLELEGN |
| Y-ATRAM | GYAGLAGLLGLEGLLGLPLGLLEGLWLGLELEGN |

### p*K*_FI_ determination

Peptides were dissolved in 10 mM sodium phosphate pH 8.0 and incubated with lipid vesicles composed of POPC. After a 1-hour incubation at room temperature, the samples were pH-corrected with 100 mM buffers (sodium acetate, MES, HEPES, or sodium phosphate). The final pH was measured after obtaining the fluorescence spectra. The molar lipid-to-peptide ratio was 200:1 with a final peptide concentration of 1 μM. The lipid-to-peptide ratio was changed to 150:1 for experiments studying the effect of salt due to the scattering effect of sodium chloride on the lipid vesicles. The final sodium chloride concentration in those experiments was 150 mM. Tryptophan fluorescence emission spectra was measured on a QuantaMaster fluorometer (Photon Technology International, Edison, NJ) with λ_ex_ = 280 nm and λ_em_ = 310-400 nm. The emission polarizer was set to 90°, and the excitation polarizer was set to 0° to reduce the light scattering effect of liposomes^*13*^. Appropriate lipid backgrounds were subtracted in all cases. Fluorescence intensities at 335 nm (F) were fitted to determine the *pK*_FI_, using:

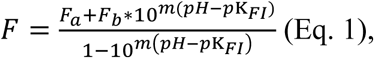

where *F*_*a*_ is the acidic baseline, *F*_*b*_ is the basic baseline, *m* is the slope of the transition, and *pK*_FI_ is the midpoint of the curve.

We then varied the composition of the lipid vesicles by gradually increasing the molar percent of PS (specifically, POPS) from 0% to 20%. The amount of PS needed to reach 50% of change in p*K*_FI_ was defined as PS_50_ and calculated according to:

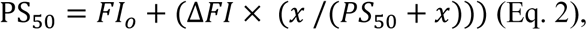

where *FI*_o_ is the p*K*_FI_ at 0% PS, *ΔFI* is the change in the p*K*_FI_, and *x* is the molar percentage of PS.

### Circular dichroism

Peptides were prepared in 10 mM sodium phosphate pH 8.0 and incubated with POPC or POPC/POPS (9/1 molar ratio) vesicles. The pH of the samples was adjusted with 100 mM sodium acetate pH 4 or 100 mM sodium phosphate pH 8 after a 1-hour incubation at room temperature. The final pH was measured after measuring the spectra. The lipid-to-peptide ratio was 200:1 with a final peptide concentration of 5 μM. Appropriate blanks were subtracted. Measurements were performed on a Jasco J-815 spectropolarimeter (Easton, MD) at room temperature. Raw data were transformed to mean residue ellipticity according to

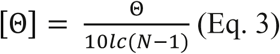

where Θ is the measured ellipticity in millidegree, *l* is the path length of the cuvette in cm, *c* is the peptide concentration in M, and *N* is the number of residues.

### Oriented circular dichroism

Lipid or lipid-peptide (50:1 molar ratio) films were resuspended in methanol and added onto two circular quartz slides (Hellma Analytics, Germany). The slides were placed under vacuum for 24 hours to evaporate the solvent. The samples on the slides were rehydrated with 100 mM sodium acetate pH 4 for 16 hours at room temperature at 96% relative humidity. The slides with the samples were placed on opposite sides of the OCD cell in a manner that would seal the cell. This ensured that the saturated K_2_SO_4_ that filled the inner cavity of the cell would remain hydrated and constantly humidify the sample throughout the experiment. The sample was recorded on a Jasco J-815 spectropolarimeter at room temperature. Eight 45° angle intervals were measured to prevent the artifacts caused by linear dichroism and then averaged for the final spectrum. Data were converted into mean residue ellipticity after lipid blanks were subtracted. The theoretical transmembrane and peripheral helix spectra was calculated as described by Wu and colleagues^*14*^.

### NBD lipid binding assay

NBD-labeled peptides were prepared in 10 mM sodium phosphate pH 8.0 and incubated with various POPC or POPC/POPS (9/1 molar ratio) concentrations at a final peptide concentration of 0.2 or 0.5 μM. The lipid-to-peptide ratios were maintained over the two peptide concentrations. After 1 hour incubation at room temperature, the pH of the samples was changed with 100 mM sodium acetate pH 4.05 or 100 mM sodium phosphate pH 7.5. Fluorescence spectra were measured on a Cytation 5 microplate reader (BioTek Instruments, Inc., Winooski, VT) with λ_ex_ = 470 nm and λ_em_ = 520-600 nm. Fluorescence intensities at 540 nm (F) were normalized to the highest value of each individual binding curve and were fitted with:

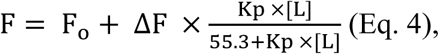

where F_o_ is the initial fluorescence intensity, ΔF is the overall change in fluorescence signal, *K*_*p*_ is the partition coefficient, [L] is the concentration of lipids, and 55.3 is the molar concentration of water.

### Sulforhodamine B leakage assay

Dequenching experiments were used to measure the release of sulforhodamine B (SRB) from POPC or POPC/POPS (9/1 molar ratio) vesicles. The dye was encapsulated into large unilamellar vesicles (LUVs) by resuspending the dried lipid film with 20 mM SRB and extruded with 200 nm filters. The SRB-LUV suspensions were purified with a PD-10 column. A consistent volume of peptide at different initial concentrations was added to the SRB-LUV suspensions to extend a range of lipid-to-peptide ratios. Samples with either 10 mM sodium phosphate pH 8 or 1.5% Triton X-100 added instead of peptide were used as controls. The pH of the samples was adjusted with 100 mM sodium acetate pH 4.05 or maintained with 100 mM sodium phosphate pH 7.5 after a 1-hour incubation at room temperature. The change in fluorescence intensity was measured after an additional 1-hour incubation on a Cytation 5 microplate reader with λ_ex_ = 550 nm and λ_em_ = 590 nm. SRB leakage was calculated with the following equation:

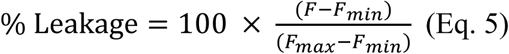

where *F, F*_*max*,_ and *F*_*min*_ are the measured fluorescence intensities of SRB-LUV suspensions with peptide, Triton X-100, and buffer, respectively. A correction was applied to account for the change in SRB fluorescence caused by Triton X-100.

### Statistical analysis

Statistical analysis on p*K*_FI_ and *K*_p_ was performed using the SPSSv25 software (IBM Analytics, Armonk, NY). Levene’s test was performed to determine the homogeneity of the variance. Multiple comparisons tests were chosen based on the homoscedasticity (Tukey) or heteroscedasticity (Dunnett T3) of the data. Comparison of the p*K*_FI_ values was performed using the Tukey test, while the *K*_p_ was compared using the Dunnett T3 test. *p* ≤ 0.05 was considered significant. The *n* values presented in the figure legends refer to biological replicates consisting of independent repetitions of the full protocols.

## Results

### ATRAM variants

The molecular basis of the interaction of ATRAM with lipid membranes is poorly understood. We performed mutational analysis to fill this void in knowledge. When ATRAM folds into the membrane forming a transmembrane (TM) helix, it does so inserting the C-terminal end (C_t_) across the membrane^*15*^. To avoid disrupting the overall membrane interaction of the peptide, we did not mutate the C_t_. Instead, we modified the N-terminus (N_t_) to create two peptide variants, named K2-ATRAM and Y-ATRAM (Table 1).

We first performed studies at pH 8, where ATRAM binds to the membrane surface, and then at pH 4, which triggers ATRAM to adopt a TM orientation^*5*^. Tryptophan fluorescence emission (Fig. 1) showed that the overall pH-responsiveness of the two new peptide variants was similar to ATRAM. This was the case in the presence of both POPC and POPC/POPS (molar ratio = 9/1) lipid vesicles. Specifically, we observed that upon a pH decrease, the fluorescence intensity increased, and the maximum of the spectra blue-shifted more than 10 nm (Fig. 1 and Table 2). These results indicate that upon acidification the single W residue (Table 1) transitioned from a polar environment to a more hydrophobic medium. The CD spectra showed that at pH 8 all peptides were mostly unstructured, both in buffer and in lipid vesicles. As expected, they all formed alpha helices in the presence of lipid vesicles at pH 4 (Fig. S1).

**Table 2.** Fluorescence spectral maxima (nm) at acidic and basic pH values.

|  | ATRAM | K-ATRAM | Y-ATRAM |
| --- | --- | --- | --- |
| **POPC acidic** | $327.3 \pm 0.7$ | $329.7 \pm 1.2$ | $324.5 \pm 2.9$ |
| **POPC basic** | $341.1 \pm 2.3$ | $342.1 \pm 2.1$ | $343.9 \pm 3.6$ |
| **POPC/POPS acidic** | $330.8 \pm 0.8$ | $320.2 \pm 1.7$ | $320.5 \pm 1.9$ |
| **POPC/POPS basic** | $350.9 \pm 1.0$ | $331.7 \pm 1.8$ | $342.7 \pm 2.3$ |

**Figure 1.**
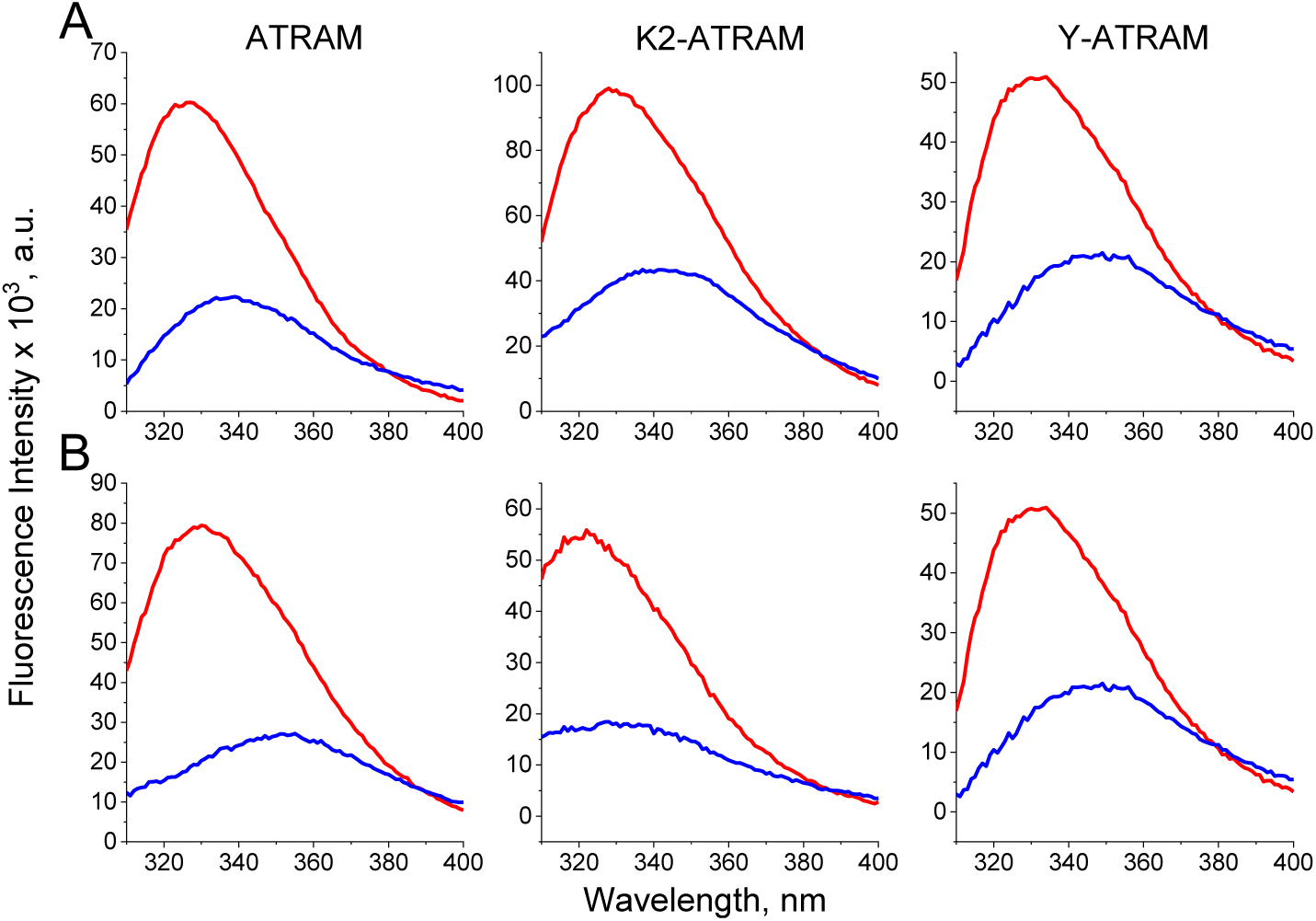
ATRAM, K2-ATRAM and Y-ATRAM interact with the lipid membrane in a pH dependent manner. Representative intrinsic tryptophan fluorescence spectra in the presence of POPC (A) and POPC/POPS (9/1) (B) vesicles at pH 4 (red) and pH 8 (blue).

Oriented circular dichroism (OCD) was performed to study the helical orientation of the peptides with respect to the plane of the bilayer at acidic pH. An α-helix laying on top of a lipid bilayer generates an OCD spectrum with two clear minima at ∼205 and ∼222 nm. However, the OCD spectrum of a TM helix has a single broad minimum at ∼225 nm. Our experimental data indicate that all peptides adopted a TM orientation at pH 4 in both lipid compositions (Fig. 2 and Fig. S2), with a small helical tilt. Theoretical oriented circular dichroism (OCD) curves in POPC vesicles are shown along the measured curves of the peptides (Fig. 2). The variation between the theoretical curves of the different peptides arises from the fact that the theoretical curves depends on the fractional helicity of the peptides^*14, 16*^. Taken together, these studies demonstrate that the mutations do not hamper pH-dependent membrane insertion.

**Figure 2.**
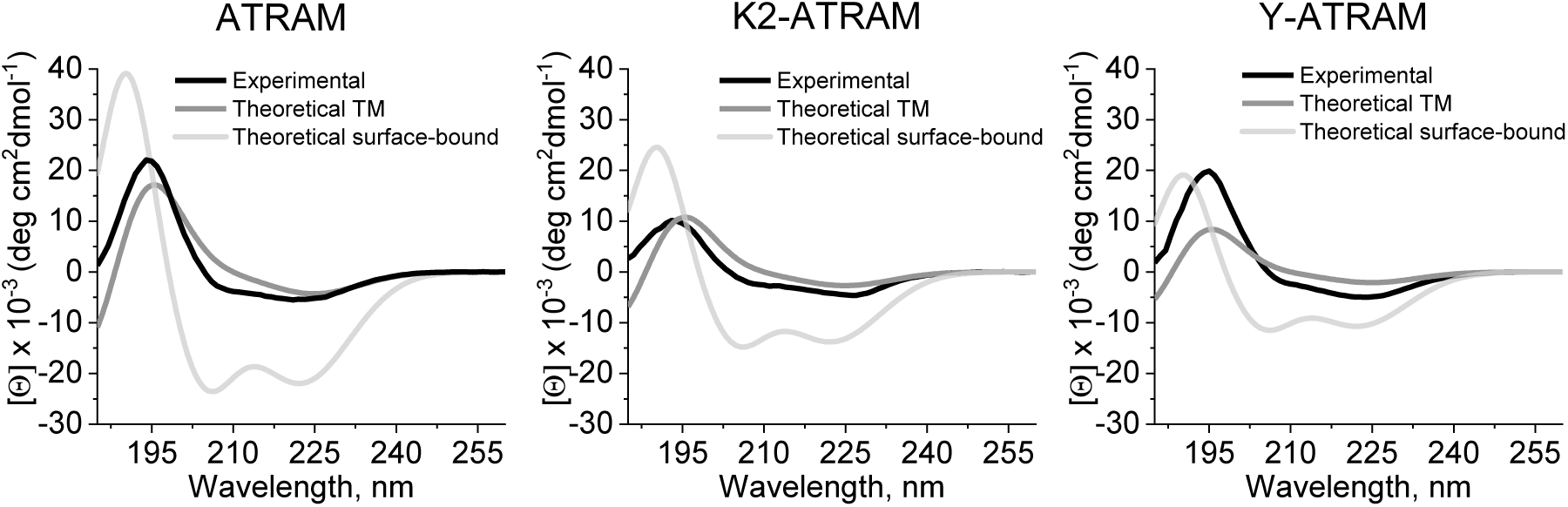
All peptides adopt a TM conformation at acidic pH. OCD spectra of the peptides (black) in hydrated stacked POPC bilayers. The grey lines correspond to theoretical spectra for a TM (dark grey) and surface-bound (light grey) α-helix.

### p*K* of membrane insertion

To determine the pH midpoint of the membrane insertion of the peptides, we followed the changes of the tryptophan fluorescence intensities as the pH decreased from pH ∼8 to pH ∼4 (Fig. 3). The pH midpoint of this sigmoidal titration is defined as the p*K*_FI_^*17*^. In all cases we observed that acidification caused sigmoidal fluorescence changes. In experiments performed with POPC vesicles, the p*K*_FI_ of ATRAM (6.20 ± 0.15) was similar to that of Y-ATRAM (6.27 ± 0.12). However, for K2-ATRAM we observed a large acidic shift, resulting in a p*K*_FI_ of 5.42 ± 0.14.

**Figure 3.**
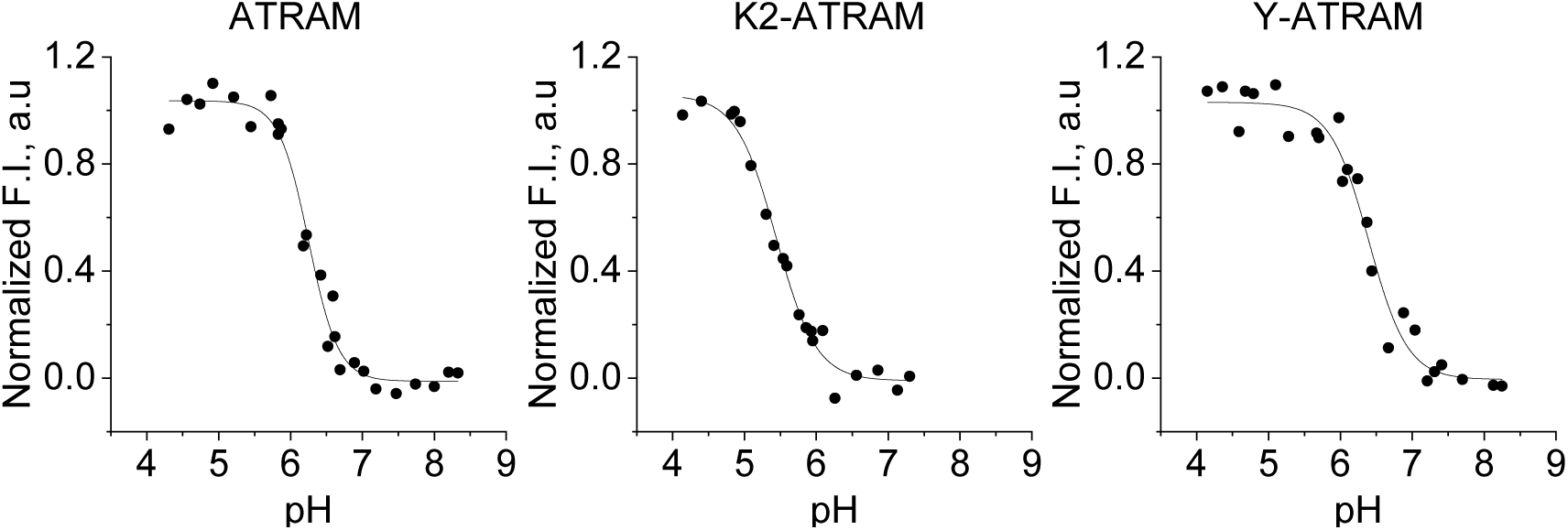
Representative pH titration curves of ATRAM and the two variants. The p*K*_FI_ of each peptide in POPC was determined as the midpoint of the change in the peptide intrinsic fluorescence intensities, obtained by fitting to Eq. 1.

We studied if similar results were observed when PS was present in the vesicles. To this end we repeated the pH titrations with seven PS levels between 0-20%, to cover the physiologically relevant range^*18*^. Intriguingly, we observed that the impact of PS in the p*K*_FI_ was markedly different between the three peptides (Fig. 4A and Fig. S3). The p*K*_FI_ of ATRAM was not significantly affected by the presence of PS in the lipid vesicles. However, there was a different, and opposite, behavior for K2-ATRAM and Y-ATRAM. Thus, PS caused a ∼0.4 pH units increase in the p*K*_FI_ for K2-ATRAM. However, PS caused a decrease in the p*K*_FI_ for YATRAM (∼0.3 pH units). The PS changes were hyperbolic, and fitting to Eq. 2 allowed determining the PS midpoint (PS_50_). The PS_50_ values were 1.9% PS for K2-ATRAM, and 0.5% PS for Y-ATRAM. These low values illustrate that the effect on the p*K*_FI_ occurred at low concentrations of PS, well within those present in most eukaryotic cells^*18*^.

**Figure 4.**
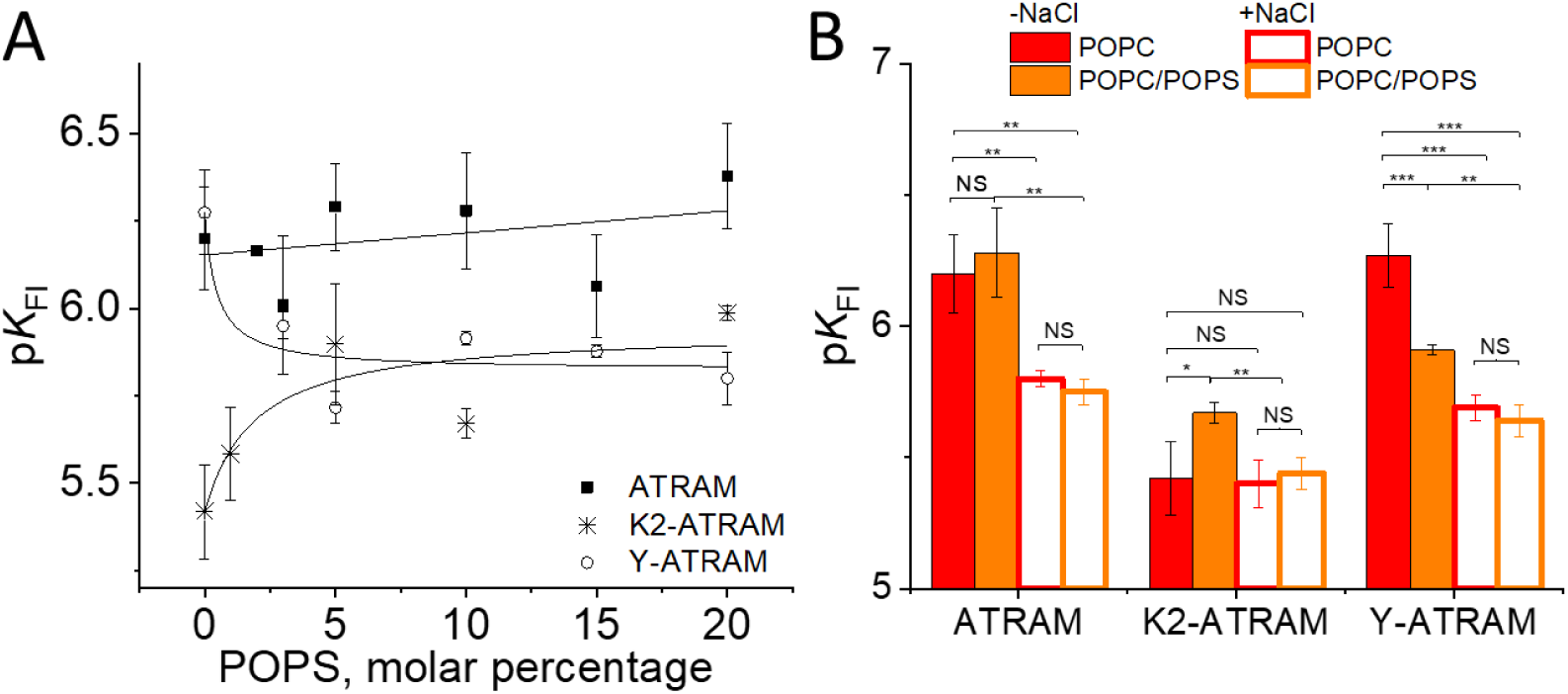
The effect of POPS and NaCl on the p*K*_FI_. (A) Increasing levels of PS effect on ATRAM (squares), Y-ATRAM (circles) and K2-ATRAM (stars). (B) Effect of the addition of 150 mM NaCl on the p*K*_FI_ in POPC or POPC/POPS (9/1). Mean pK_FI_ values from panel A plus additional data obtained in the presence of NaCl are shown ± S.D. *n* = 3-6, \**p*<0.05, \*\**p*<0.01,\*\*\**p*<0.001, NS = no significance.

The effect of PS on the p*K*_FI_ of Y-ATRAM is similar to the effect we had previously reported with pHLIP, a different pH-sensitive membrane peptide^*19*^. For pHLIP, the p*K*_FI_ decrease was due to electrostatic repulsion between the negative charges in PS and pHLIP, which was screened in the presence of NaCl. To study whether electrostatics also controlled the membrane insertion of the three ATRAM peptides, the p*K*_FI_ was also determined in the presence of 150 mM NaCl (Fig. 4B and Fig. S3). For ATRAM, the p*K*_FI_ of both lipid conditions decreased compared to the values obtained in the absence of NaCl. Similarly, a p*K*_FI_ decrease was observed for Y-ATRAM for both lipid compositions. In the presence of NaCl both peptides showed similar p*K*_FI_ in POPC and POPC/POPS.

A more nuanced scenario was observed for K2-ATRAM, as NaCl decreased p*K*_FI_ only in POPC/POPS, but not for POPC liposomes (Fig. 4B). Again, the difference in p*K*_FI_ between POPC and POPC/POPS vesicles was lost in the presence of NaCl. An additional feature of K2-ATRAM is that the p*K*_FI_ in the presence of NaCl (∼5.4) was lower than for ATRAM and YATRAM, with a value in both cases of ∼5.7. These results indicate that fundamental differences exist in how K2-ATRAM and the two other peptides interact with lipid membranes. Due to lipid light scattering, titrations in the presence of NaCl were performed with a lipid-to-peptide ratio of 150:1, instead of 200:1. We performed control experiments that showed that in the presence of 150 mM NaCl there were no significant differences in the p*K*_FI_ of ATRAM at 150:1 and 200:1 POPC-to-peptide ratios, (Fig. S4). To unravel the causes for the different effect of PS on the three ATRAM peptides, we performed additional studies.

### Partition coefficient

We have previously reported that the POPC affinity of ATRAM is higher at acidic pH than neutral pH, as a consequence of the increased hydrophobicity resulting from the protonation of glutamic acids^*15*^. We obtained similar results were also observed for all ATRAM variants in POPC (Fig. 5). Interestingly, at acidic pH the effect of POPS on the partition coefficient mirrors the trends seen in the p*K*_FI_ (Fig. 4B). Thus, at pH 4 the presence of POPS did not affect the *K*_p_ for ATRAM (Fig. 5), but it increased for K2-ATRAM (as it increased p*K*_FI_) and decreased for Y-ATRAM (as it decreased p*K*_FI_). Previously, the binding of ATRAM to the lipid membrane was considered non-ideal, as the partition coefficient depended on the peptide concentration, and thus the *K*_p_ we report is in reality an apparent *K*_p_^*15*^. Here we show that this was also true for all the variants in both lipid composition, as when we repeated the binding assay at a lower peptide concentration, the *K*_p_ values were significantly higher (Figs. S5 and S6). However, similar trends were observed in both cases.

**Figure 5.**
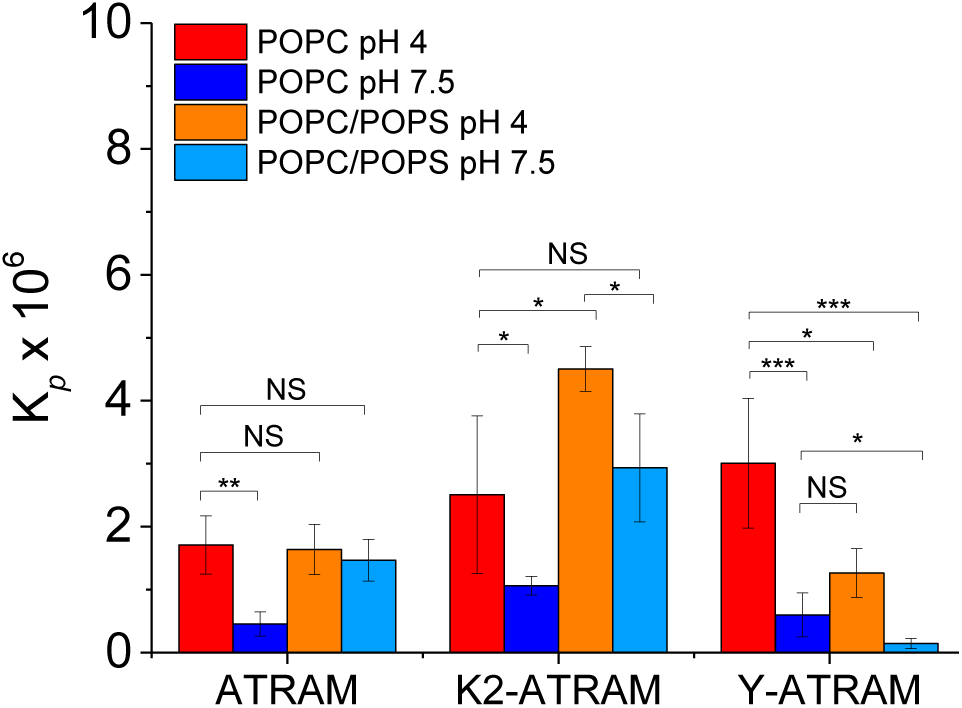
Apparent *K*_p_ of ATRAM peptides in POPC and POPC/POPS (9/1). Mean values are shown ± S.D. *n* = 3-4, \**p*<0.05, \*\**p*<0.01, \*\*\**p*<0.001, NS = no significance.

### Membrane leakage

ATRAM oligomerizes on the membrane of POPC vesicles^*5*^. Peptide oligomerization is often linked to membrane disruption^*20*^. Next, to gain insights into their oligomerization, we studied the effect of the peptides on leakage of encapsulated sulforhodamine B (SRB). We performed a SRB leakage assay in POPC and POPC/POPS (molar ratio = 9/1) vesicles (Fig. 6), where SRB de-quenching occurs as it leaks out of the vesicles. Overall, similar leakage results in each lipid composition were obtained for the three peptides. These results suggest that no large oligomerization differences might exist between the three variants.

**Figure 6.**
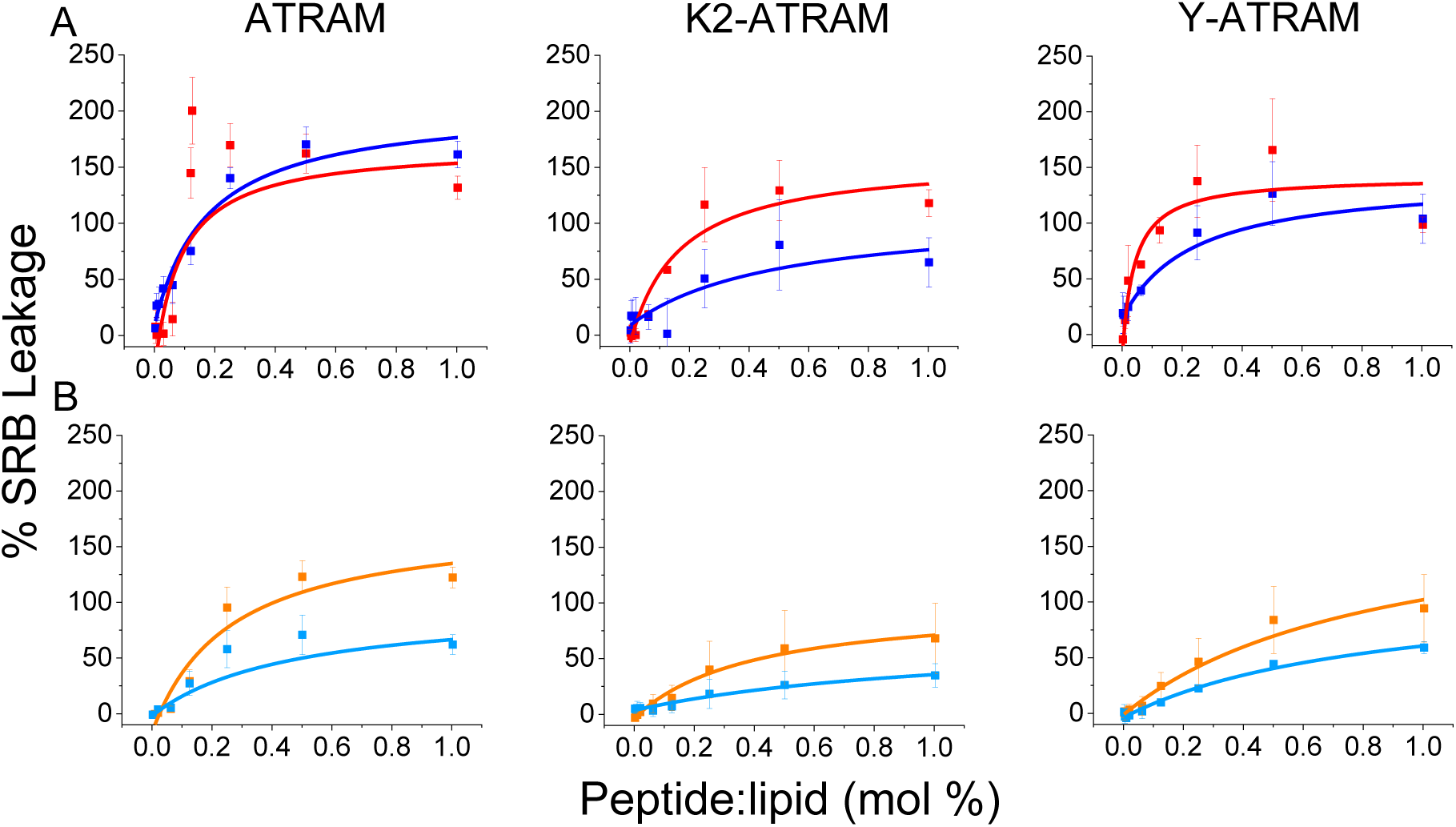
Membrane leakage studies. De-quenching of sulforhodamine B (SRB) as it is released from vesicles of POPC (A) or POPC/POPS (9/1) (B). Fluorescence was monitored after the addition of the three peptides (0.0025-1 mol %) at pH 7.5 (blue) and pH 4 (red/orange). Complete leakage was achieved on addition of Triton X-100. Fluorescence was corrected, since Triton X-100 altered the fluorescence intensity of SRB in solution. Deviations from a maximal leakage of 100% are attributed to errors in this correction. Mean values are shown ± S.D. (*n* = 4).

## Discussion

PS the is the most abundant negatively charged lipid in the plasma membrane of eukaryotic cells^*18*^. PS plays multiple regulatory roles on membrane proteins, that result in changes in activity^*21*^, polymerization^*22*^ and clustering^*23*^, as well as driving membrane interaction of soluble peripheral membrane proteins^*24*^, and myristoylated proteins^*25*^. PS is distributed asymmetrically in the plasma membrane of human cells, with high levels in the cytoplasmic leaflet, and low levels present in the extracellular leaflet^*26, 27*^. PS asymmetry, however, can be lost in cancer cells, where PS is increasingly exposed at the cell surface^*28-30*^. This provides the opportunity of targeting PS to direct selective drug delivery to cancer cells.

In this work, we studied the effect on ATRAM of the presence of POPS in the membrane. We designed two new ATRAM variants to study the effect of the properties of the peptide N_t_. Fluorescence and circular dichroism confirmed that the variants maintained pH-sensitive properties. Furthermore, OCD experiments in POPC showed that all peptides acquired a TM conformation at low pH. The OCD spectra of the peptides in POPC/POPS had a lower signal than those in POPC (Fig. S2). A possible explanation is that that the negative charges of PS caused bilayer repulsion^*31*^, making it harder for lipid bilayers to stack on top of each other on the quartz slide, artifactually reducing the overall signal.

We determined that the p*K*_FI_ of ATRAM in POPC vesicles was 6.2. However, the previously published insertion p*K* value of ATRAM in POPC was 6.5^*5*^. This latter value was determined by following the shift of the fluorescence spectral maximum over a pH range, while the new value was determined by following instead changes in intensity. It has previously been shown for the pHLIP peptide that different methods of spectral analysis result in changes in p*K* values, as each method reported different membrane insertion intermediates^*32*^. Previous stopped-flow fluorescence data showed that ATRAM populates at least three intermediates when transitioning from the peripheral state to the TM conformation^*15*^. This result suggests that the observed differences in p*K* values of ATRAM might be associated with the preferential detection of different conformational intermediates.

In POPC vesicles, we observed a decreased p*K*_*FI*_ for K2-ATRAM compared to ATRAM and Y-ATRAM (Fig. 4). This result indicates that the addition of the two basic residues resulted in more protons being required for the peptide to insert. Studies in the presence of NaCl revealed that ATRAM and Y-ATRAM electrostatically interact with POPC, but this was not the case with K2-ATRAM, since there was no change in p*K*_FI_ with the addition of NaCl. As tumor cells expose PS on the outer leaflet of their cell membranes^*33*^, we studied the effect of PS on the membrane interaction of the peptides. While PS did not affect the p*K*_*FI*_ of ATRAM, it affected the two variants in an opposite fashion. For K2-ATRAM, the observed increase in p*K*_FI_ is expected to result from favorable interactions with the negative charge of PS. As with ATRAM and Y-ATRAM, the addition of NaCl resulted in a decrease in p*K*_FI_ for K2-ATRAM, indicating that the interaction with PS has an electrostatic component for all peptides. The NaCl experiments were also performed at a lower lipid-to-peptide ratio (150:1 *vs.* 200:1). While the p*K*_FI_ did not differ for these two ratios (Fig. S4), we cannot rule out that any effects from working at non-saturating conditions at pH 7.5, particularly for Y-ATRAM (Fig. S6).

The complexity in the effect of PS and NaCl on the peptides suggests that the interaction is not controlled by a simple electrostatic process ^*19*^. For pHLIP, it is proposed that the decrease in membrane insertion p*K* caused by PS results from the protonatable residues being more hydrated ^*19*^. This would result from the peptide having a shallower position on the membrane surface due to the presence of negatively charged lipids. As a result, there would be enhanced exposition to the aqueous environment of the sidechains of the protonatable (Asp and Glu) residues. The spectral maximum of the C_t_ Trp is sensitive to hydration and thus reports the degree of membrane embedding, with a lower spectral maximum indicating deeper insertion into the hydrophobic bilayer (Table 2). In ATRAM, the single Trp is surrounded by three of the four Glu residues. However, we did not observe a good correlation between the Trp hydration and the p*K*_FI_. Then, we hypothesize that the remaining Glu residue, away from the Trp at position 12 (E12), controls the insertion p*K*_FI_ changes.

Integration of the data of Table 2 and Fig. 4A allows us to propose a working model for the interaction of the peptides with the surface of the membrane (Fig. 7). In ATRAM, the residues W26 and E12 are similarly immersed on the lipid polar head group both in POPC and POPC/POPS vesicles, resulting in a similar p*K*_FI_. In K2-ATRAM, in POPC the N_t_, including E12, is away from the membrane, resulting in a lower p*K*_FI_, which is closer to ∼4.0-4.5, the value typically observed for Glu in solution^*34*^. However, when POPS is present in the membrane, an attractive interaction occurs between the negative charges of PS and the positively charged lysines. As a result, the N_t_ approaches the membrane, altering the polarity of the environment of E12 to increase the p*K* of this residue. In the case of Y-ATRAM, the cause of the effect of PS on the p*K*_FI_ is less clear. However, we hypothesize an involvement of the polar nature of the Tyr ring, which provides a surface of negative electrostatic potential that could cause electrostatic repulsion with the PS headgroup^*35*^. As a result, E12 would more exposed to the aqueous environment, which is expected to decrease the insertion p*K*_FI_ in the presence of PS. Fig. 7 only shows the surface-bound state, as we propose that the TM state is similar in all cases. Interestingly, we observed a correlation between the membrane affinity at neutral pH (Fig. 5, K*p*) and the pH-responsiveness of insertion (Fig. 4, p*K*_FI_). The increase in affinity in the presence of POPS observed in K2-ATRAM was probably due to favorable electrostatic interactions of the positively charged Lys and the negatively charged PS, while the opposite was observed with Y-ATRAM. This highlights the importance for peripherally binding peptides and proteins of considering the equilibrium between surface-bound and soluble states.

**Figure 7.**
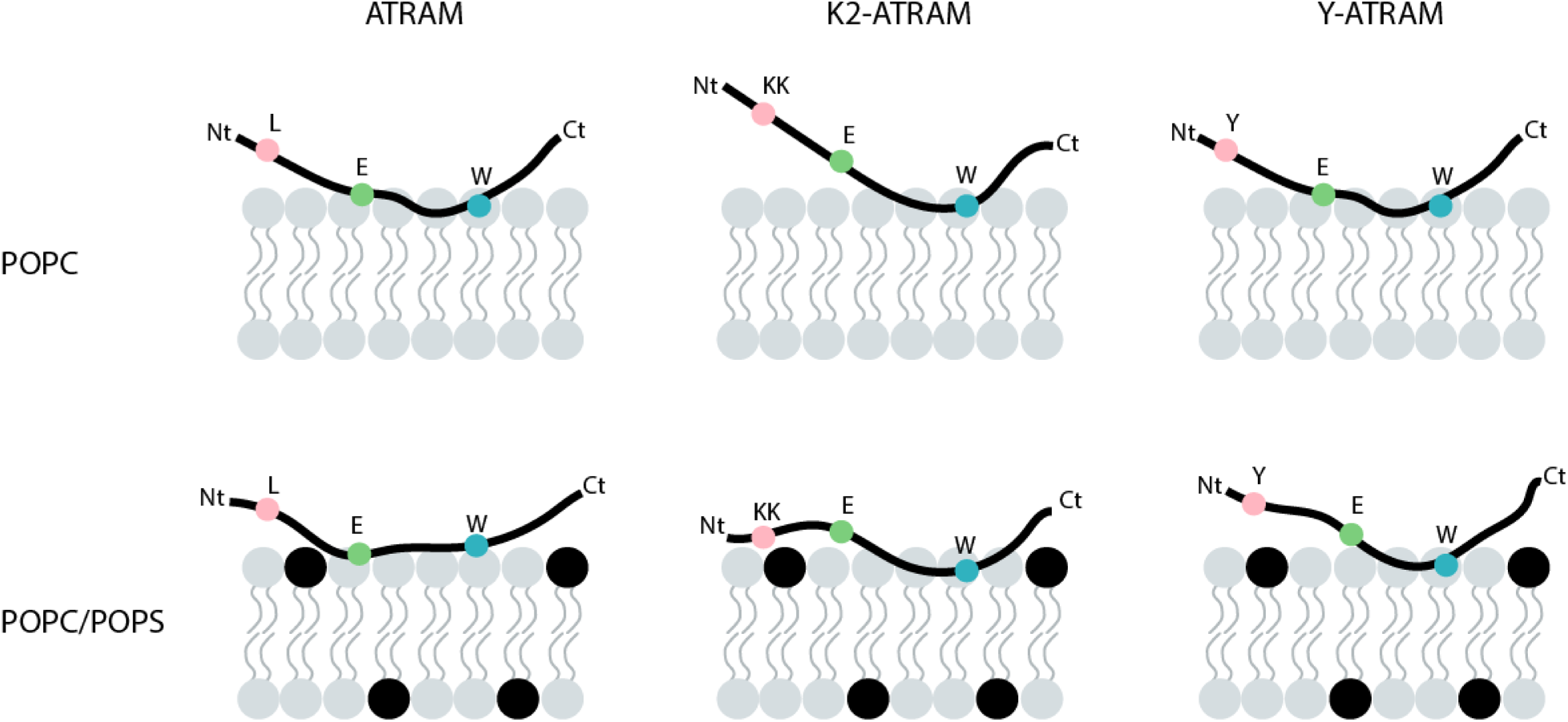
Schematic of the surface-bound state of the ATRAM peptides. The proposed locations of E12 and W26 residues are shown in green and blue, respectively, while the modified residues at the N-terminal are marked in pink. The two lysines are represented as a single sphere. The headgroup of PS is highlighted in black.

We observed significant leakage for all peptides both in POPC and POPC/POPS (9/1), suggesting that the ATRAM variants had the ability to oligomerize into membranes. However, we have previously shown that any membrane disruptions caused by ATRAM are not significant enough to cause cell death in cultured cells^*5*^. SRB is a small molecule (MW = 558.6 Da), and as a result SRB leakage assays might report on relatively small perturbations to the lipid membrane that do not occur in the more complex and dynamic plasma membrane. Furthermore, we have previously reported that ATRAM did not induce significant membrane leakage of calcein^*5, 15*^. This is consistent with a previous report showing that SRB displays enhanced leakage, suggesting that SRB assays might be over-reporting true membrane leakage^*36*^.

This work shows that the sequence opposite to the inserting end can control the insertion of a membrane peptide. This indicates that for ATRAM, as well as for similar peptides such as TYPE7^*37*^ and pHLIP^*38*^, the N_t_ is a promising region for modifications that improve tissue targeting or pharmacokinetics. In our case, the p*K* of insertion of the three peptides still remained more acidic than the pH of the extracellular matrix of tumor cells (pH 6.4–7.0) ^*2, 39*^. However, it is important to consider that the local pH at the membrane surface is significantly more acidic that the bulk microenvironment pH^*40, 41*^. This suggests that the pH-triggered membrane insertion of ATRAM peptides could provide specificity for tumor targeting. Indeed, it has been recently reported that ATRAM efficiently targets tumors and is able to translocate drug-like molecules across the plasma membrane of cancer cells^*6*^. Our results point towards the intriguing possibility that the exposure of PS in cancer cells could be used as an additional source of tumor specificity^*42*^. Indeed, for K2-ATRAM, the presence of small amounts of PS rapidly increases the p*K*_FI_, which is expected to increase cancer cell targeting. This might be the basis for K2-ATRAM potentially displaying an increased specificity for tumor cells. The opposite would be expected for the Y-ATRAM variant. We propose that our findings can be used to achieve improved design of peptides that target the membrane of cancer cells, particularly in the more acidic aggressive solid tumors.

## Author Contribution

VPN performed research, analyzed data and wrote the paper, ACD performed research and analyzed data, and FNB designed research, analyzed data and wrote the paper.

## Acknowledgements

We are thankful to Justin M. Westerfield and Yujie Ye for insightful comments on the manuscript. This work was supported by grant R01GM120642 to F.N.B.

